# Single-Cell RNAseq of Out-of-Thaw Mesenchymal Stromal Cells Shows Striking Tissue-of-Origin Differences and Inter-donor Cell-Cycle Variations

**DOI:** 10.1101/2020.09.10.290155

**Authors:** Camila Medrano-Trochez, Paramita Chatterjee, Pallab Pradhan, Molly E Ogle, Edward A Botchwey, Joanne Kurtzberg, Carolyn Yeago, Greg Gibson, Krishnendu Roy

**Affiliations:** School of Biology, Georgia Institute of Technology, Atlanta, GA, 30332, USA; Marcus Center for Therapeutic Cell Characterization and Manufacturing, Institute for Bioengineering and Bioscience, Georgia Institute of Technology, Atlanta, GA, 30332, USA; Department of Biomedical Engineering, Georgia Institute of Technology, Atlanta, GA, 30332, USA; Marcus Center for Cellular Cures, Duke University School of Medicine, Durham, NC, 27705, USA

## Abstract

Mesenchymal stromal cells (MSCs) from a variety of tissue sources are widely investigated in clinical trials, and the MSCs are often administered immediately after thawing the cryopreserved product. While previous reports have examined the transcriptome of freshly-cultured MSCs from some tissues, little is known about the single-cell transcriptomic profiles of out-of-thaw MSCs from different tissue sources. Such understanding could help determine which tissue origins and delivery methods are best suited for specific indications. Here, we characterized cryopreserved MSCs, immediately post-thaw, from bone marrow (BM) and cord tissue (CT), using single-cell RNA sequencing (scRNA-seq). We show that out-of-thaw BM-vs. CT-MSCs have significant differences in gene expression. Gene-set enrichment analyses implied divergent functional potential. In addition, we show that MSC-batches can vary significantly in cell cycle status, suggesting different proliferative vs. immunomodulatory potentials. Our results provide a comprehensive single-cell transcriptomic landscape of clinically and industrially relevant MSC products.

**Highlights:**

- Single cell gene expression comparison between Bone-marrow derived MSCs and Cord-tissue derived MSCs
- Donor effects and cell heterogeneity on tissue-specific MSC gene expression
- Single Cell Pooling Enhances Differential Expression Analysis for Bone marrow and Cord tissue MSC samples
- Gene ontology reveals tissue specific unique molecular function and pathways

## Introduction

Mesenchymal Stromal Cells (MSCs), often referred to as Mesenchymal Stem Cells or Signaling Cells, are cells isolated from various tissues that have shown multipotent, regenerative, and immunomodulatory capacities in vitro. These cells, from a variety of tissue-sources, are being evaluated for therapeutic interventions, especially across a variety of inflammatory and immune conditions (1, 2). Numerous clinical trials have focused on the use of MSCs as a cell therapy for various diseases with unmet medical challenges, including graft-vs-host disease, osteoarthritis, autism, acute respiratory distress syndrome (ARDS), autoimmune diseases, and even COVID-19. A ClinicalTrial.gov search (date: August 7, 2020) with the keyword: MSC as other terms shows 4,044 studies that are either recruiting, not yet recruiting, enrolling by invitation, and active but not recruiting. MSCs are also widely used in developing engineered tissues ex vivo (3-5). Several are also working on developing MSC-based therapies and others are developing reagents and large-scale cell banks for eventual clinical use.

Despite such widespread interest in academia, clinical trials, and in industry, the characteristics of MSCs that are most correlative to their specific in vivo function remain unknown. MSCs can be isolated from various tissues, such as bone marrow, umbilical cord, placental, and adipose tissue, which introduces tissue-dependent variability between MSC-based cell products that may also differ according to donor. Furthermore, manufacturing processes vary between sites (both clinical and commercial), leading to process-dependent variability. These sources of variabilities across the MSC field confound the ability to compare clinical trial results and have contributed to a lack of conclusive historical data to support their potential for clinical use (6).

The International Society for Cell and Gene Therapy (ISCT) standards identify MSCs based on the expression status of a panel of specific surface markers, their ability to adhere to plastic, and their ex vivo tri-lineage differentiation potential to adipocytes, osteoblasts and chondroblasts (7, 8). However, these minimal MSC identification and functional criteria, especially surface marker expression, often do not correlate with their regenerative or immunomodulatory functions (9). Moreover, the proportion of “stromal” like progenitor cells that have high regenerative capability varies across MSC donors (10).

Therefore, there is an obvious need for deep phenotypic characterization of MSCs to compare heterogeneity as a function of tissue-of-origin as well as donor, and to identify potential phenotypic signatures that can be eventually used as predictive biomarkers or critical quality attributes (CQAs) for MSC-based products, and for understanding their putative Mechanisms of Action (MoAs).

Recently, single-cell RNA sequencing (scRNA-seq) has emerged as one of the next generation cell characterization techniques that can be used to gain deeper insight into gene transcriptional signatures at the single-cell level (11, 12). scRNA-seq enables the examination of genomes or transcriptomes of individual cells, providing a high-resolution view of cell-to-cell variation or heterogeneity within a population. Moreover, this technique can be used to explore the distinct biology of individual cells and to understand temporal cellular processes and functions, such as differentiation, proliferation, and immune response potential (13). scRNA-seq has been previously used to characterize hematopoietic differentiation (14-16) and immune cell subsets (17), including dendritic cells, monocytes (18), and innate lymphoid cells (19). A handful of reports have also used scRNA-seq to characterize differential gene expression in freshly-prepared MSCs from umbilical cord (20), adipose tissue (8), Wharton’s jelly (21) and bone marrow (22, 23).

In many clinical trial settings for allogeneic MSC-based off-the-shelf cellular therapies, to circumvent logistical and manufacturing challenges, MSC products are used post-thaw (directly from a frozen vial), rather than fresh (without freezing after culture), or culture-rescued (re-cultured after thawing) (2, 24). Since out-of-thaw MSC products could have different metabolic and functional characteristics from their fresh counterparts (24), phenotypic and functional characterization directly on the out-of-thaw MSC product is necessary to be able to find correlative attributes between their in vitro cell characteristics and corresponding clinical or pre-clinical efficacy. A comprehensive characterization of out-of-thaw MSC product from different tissue sources and donors at single-cell level may provide information on potential critical quality attributes (CQAs) and Mechanisms of Action (MoA) of the cells, and can be used to select MSC donor and/or sources for disease specific cell therapies.

In this study, we performed scRNA-seq using the drop-seq method (11, 25) to compare single-cell transcriptome profiles between commercially available bone marrow-derived MSCs (BM-MSCs) from six donors from RoosterBio Inc. (Frederick MD), and umbilical cord-tissue derived MSCs (UCT-MSCs) from four donors provided by Duke University. We characterized a total of 13 out-of-thaw samples from these ten MSC donors. Specifically, we assessed differences between individual donors as well as differences between MSC tissue sources. To overcome issues with zero counts that complicated differential expression analysis and to provide flexibility in normalization, we also introduce a new analytical framework, scPool, in which similar cells from the same donor are pooled into pseudo-cells.

## Results

### Donor effects on Bone Marrow-Derived MSC Gene Expression

A total of seven bone marrow-derived MSC (BM-MSC) samples from six donors were thawed and processed for scRNA-seq analysis (Tables 1 and 2). First, we compared two of the BM-MSC samples between pre-freeze and post-thaw conditions to understand freeze-thaw effects on MSC gene expression. Our analysis indicated a shift in the genetic profile between pre-freeze and post-thaw conditions (Figure S1). Pre-freeze samples showed significant overexpression of 1,743 genes relative to post-thaw samples at the 5% false discovery rate (FDR) threshold, while 310 genes were significantly overexpressed in the post-thaw samples compared to the pre-freeze samples. Some of the pathways overexpressed in the pre-freeze samples are cytokine signaling (*FOS, MMP2, TLN1, FOSB*), cell proliferation and cell adhesion (*ZYX, ITGA5, CLIC1* etc.), while the pathways over-expressed in the frozen, post-thaw samples are carbohydrate interconversions (*UGP2*), cholesterol/Steroid biosynthesis and regulation of apoptosis (*PSMA2, PSMB1*). Having established that the freeze-thaw process imparts substantial changes in gene expression profiles of MSCs, we focused our analyses on post-thaw MSC products from BM and CT origins, since these are being widely used in numerous clinical trials.

**Table 1.** Number of cells, average number of genes and average number of UMI per cell per BM-MSC sample.

| Samples | Lab | Number of Cells | Number of Reads | Mean Reads per cell | Median genes per cell | Number of cells post filtering | Sex | Age | Average nGenes per cell | Average nUMI per cell |
| --- | --- | --- | --- | --- | --- | --- | --- | --- | --- | --- |
| BM1 | A | 293 | 29,671,530 | 15,469 | 2,218 | 293 | Male | 25 | 2,693 | 7,581 |
| BM2a | A | 310 | 29,943,109 | 46,467 | 3,855 | 252 | Male | 21 | 3,714 | 12,242 |
| BM2b | B | 199 | 41,511,093 | 39,417 | 4,686 | 199 | Male | 21 | 4,395 | 15,506 |
| BM3 | B | 590 | 35,842,779 | 26,580 | 4,272 | 452 | Male | 22 | 4,193 | 13,415 |
| BM4 | B | 557 | 25,262,984 | 17,668 | 3,803 | 542 | Female | 26 | 3,430 | 9,718 |
| BM5 | B | 149 | 18,160,926 | 32,928 | 4,201 | 122 | Male | 31-45 | 4,057 | 13,591 |
| BM6 | B | 317 | 21,841,113 | 29,393 | 4,554 | 275 | Male | 18-30 | 4,367 | 14,385 |

**Table 2.** Number of cells, average number of genes and average number of UMI per cell per UCT-MSC sample.

| Samples | Lab | Number of Cells | Number of Reads | Mean Reads per cell | Median genes per cell | Number of cells post filtering | Sex | Age | Average nGenes per cell | Average nUMI per cell |
| --- | --- | --- | --- | --- | --- | --- | --- | --- | --- | --- |
| UCT1a | C | 161 | 16,255,153 | 19,563 | 2,472 | 161 | Male | -- | 2,808 | 8,458 |
| UCT1b | C | 251 | 31,333,645 | 38,809 | 3,840 | 251 | Male | -- | 3,667 | 12,075 |
| UCT1c | C | 251 | 31,333,645 | 38,809 | 3,840 | 251 | Male | -- | 2,477 | 7,321 |
| UCT2 | C | 692 | 59,783,150 | 25,248 | 2,979 | 547 | Male | -- | 3,053 | 9,381 |
| UCT3 | C | 397 | 43,757,708 | 26,241 | 2,724 | 396 | Male | -- | 2,720 | 9,262 |
| UCT4 | C | 284 | 20,167,531 | 29,162 | 3,717 | 259 | Male | -- | 3,616 | 11,711 |

Table 1 summarizes the sample data for all BM samples. Samples BM1 and BM2a were processed in laboratory A, while samples BM2b, BM3, BM4, BM5 and BM6 were processed in laboratory B (Table 1). An average of 305 cells were profiled per sample, with an average read depth of 29,703, representing 12,348 UMI (Unique Molecular Identifier) and 3,836 expressed genes per cell (Table 1). The profiles were clustered with both Seurat(26) and SC3 pipelines(38). Since the latter is optimized for relatively small experimental designs, we present the results of SC3, but note that similar findings were obtained with Seurat (Figure S2). Before characterizing differential expression among samples, we confirmed that the MSC-identity markers established by the ISCT, namely *NT5E (*CD73), *THY1* (CD90) and *ENG* (CD105) were detected in the majority of cells in the bone marrow and umbilical cord tissue derived MSCs (Figure S3). Furthermore, *CD34, CD14, CD19* and *PECAM1* - all markers of hematopoietic or lymphoid lineages, were absent.

Projecting each cell against the first two Principal Components (PC) of gene expression, three clusters of single cell profiles were observed (Figure 1A). PC1 is highly negatively correlated with the total UMI count per cell. Accordingly, the smallest cluster located to the right, consists of 17% of the cells all of which had low UMI counts, typically fewer than 1,000 detected genes, and low ribosomal protein transcript counts (Figure 1B). These low UMI counts cells were more prevalent in two donors studied - one from each of the two laboratories (Lab A and Lab B; BM1 and BM4, respectively: Figure 1C), suggesting the low UMI count may not be related to the lab in which they were manufactured. It is not clear whether the unusual profile of these cells is a technical artefact, or has a biological basis, but they appear to be of low quality and were excluded from all subsequent analyses.

**Figure 1.**
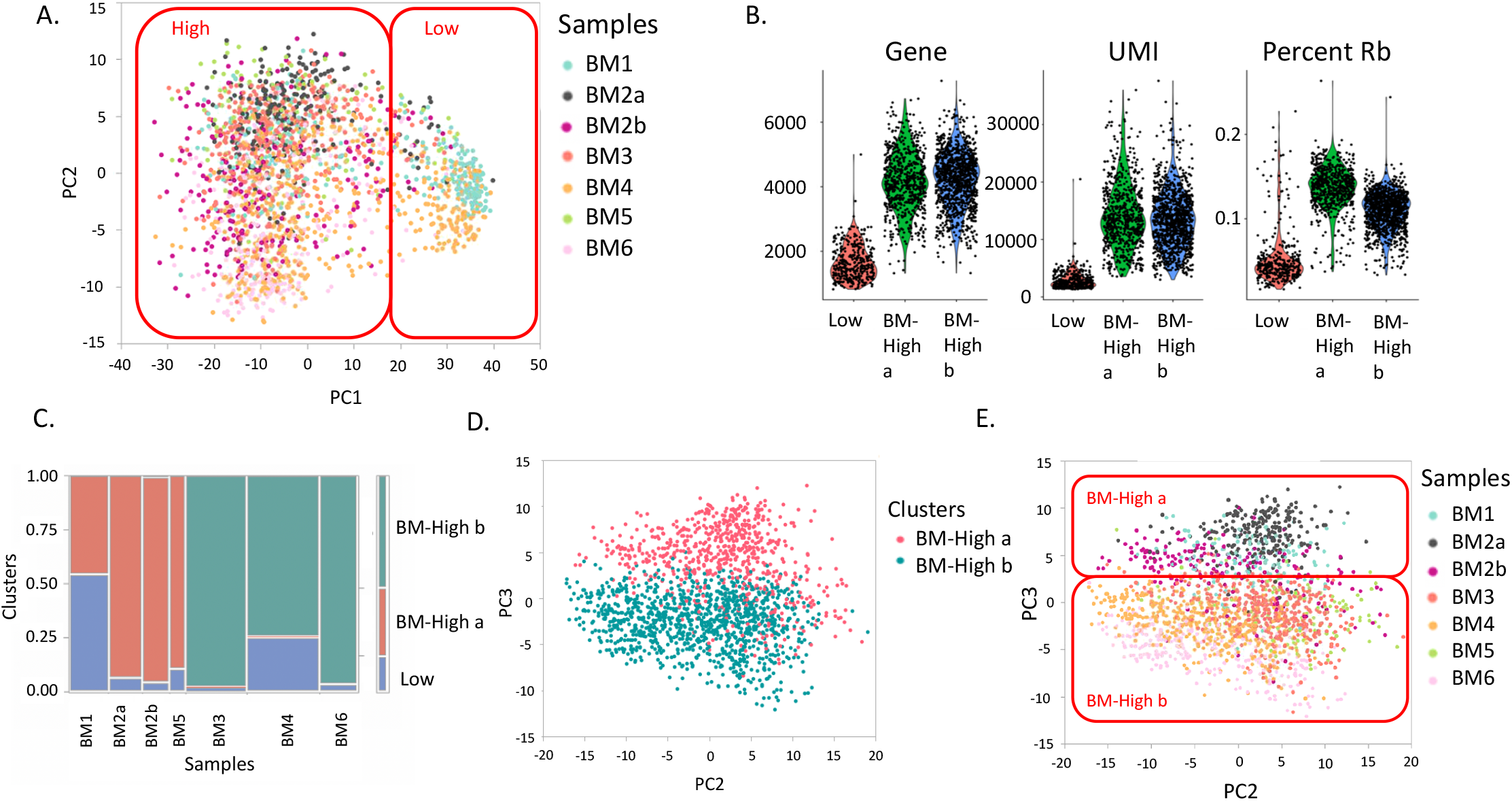
Clusters of BM-MSC transcriptome profiles. (A) The first two Principal Components of gene expression identify two broad clusters of cells, which are colored by sample: Cluster BM-Low, which correspond low UMI count cells and cluster BM-High, with high UMI count cells. (B) Violin plots show the density of the number of Genes, UMI, and Ribosomal Protein transcripts (RP) per cell. (C) Association of cells with clusters. The width of each column is proportional to the number of cells in the indicated sample, and the color of each box corresponds to cells in cluster BM-Low (blue), BM-High_a (red) or BM-High_b (green). (D) Within cluster High, SC3 identifies two clusters of cells, which separate along PC3 as indicated by the red and blue points. (E) Shading of cells by sample confirms that cells from each donor belong to one of the two sub-clusters, although with subtle separation associated with PC3.

Two clusters identified by SC3 in the remaining high-quality datasets largely differentiate along PC3 (Figure 1D). Three of the five samples expanded in laboratory B (BM3, BM4 and BM6) were predominantly found in cluster BM-High_b; the other two samples along with both of the samples expanded in laboratory A (BM1, BM2a, BM2b and BM5) were predominantly found in cluster BM-High_a (Figure 1E). Both samples from the donor whose cells were cultured in each of the laboratories are in cluster BM-High a (BM2a and BM2b), suggesting that the difference is more likely to be donor-related than due to a laboratory or technical effect. Nevertheless, Figure 1E shows that even between the two laboratories, the cells from this donor tend to separate along PC3.

### Single Cell Pooling Enhances Differential Expression Analysis for Bone Marrow-MSC samples

Although the above analysis follows current standard practice, it is nevertheless subject to the caveat that for a large percentage of the genes, transcripts are only observed in fewer than half of the cells, and consequently the differential expression analysis is mostly based on presence-versus-absence. In order to lend robustness to the conclusions, we elected to analyze differential expression with an alternative strategy that accounts for the impact of a high proportion of drop-outs in scRNA-seq. Our method, “scPool”, is based on pooling of cells of the same type within samples, which ensures that a high proportion of genes are represented by an approximately normal count distribution. It also allows for fitting of normalization strategies initially developed for bulk microarray or RNA-seq analysis.

After normalizing gene expression values to counts per 10,000 UMI, differential expression analysis between clusters BM-High_a and BM-High_b was performed in EdgeR with donor as a random covariate, yielding 4,624 genes at a FDR of 5%. There were 2,230 genes upregulated in cluster 2a, and 2,394 upregulated in cluster 2b (Figure 2A). Gene ontology analysis detected strong enrichment for multiple pathways involved in cell cycle regulation in cluster BM-High_b (Figure 2B). By contrast, cluster BM-High_a showed upregulation of multiple pathways related to immune signaling and other processes expected to be characteristic of functional MSCs (Figure 2B). On the basis of the cell cycle gene expression, cells in cluster BM-High_b may be preparing for or undergoing cell cycle division, whereas the cluster BM-High_a MSCs may be more likely to be in G0 phase. Alternately the two populations may simply express cell cycle related genes at different levels, without this reflecting cell cycle stages.

**Figure 2.**
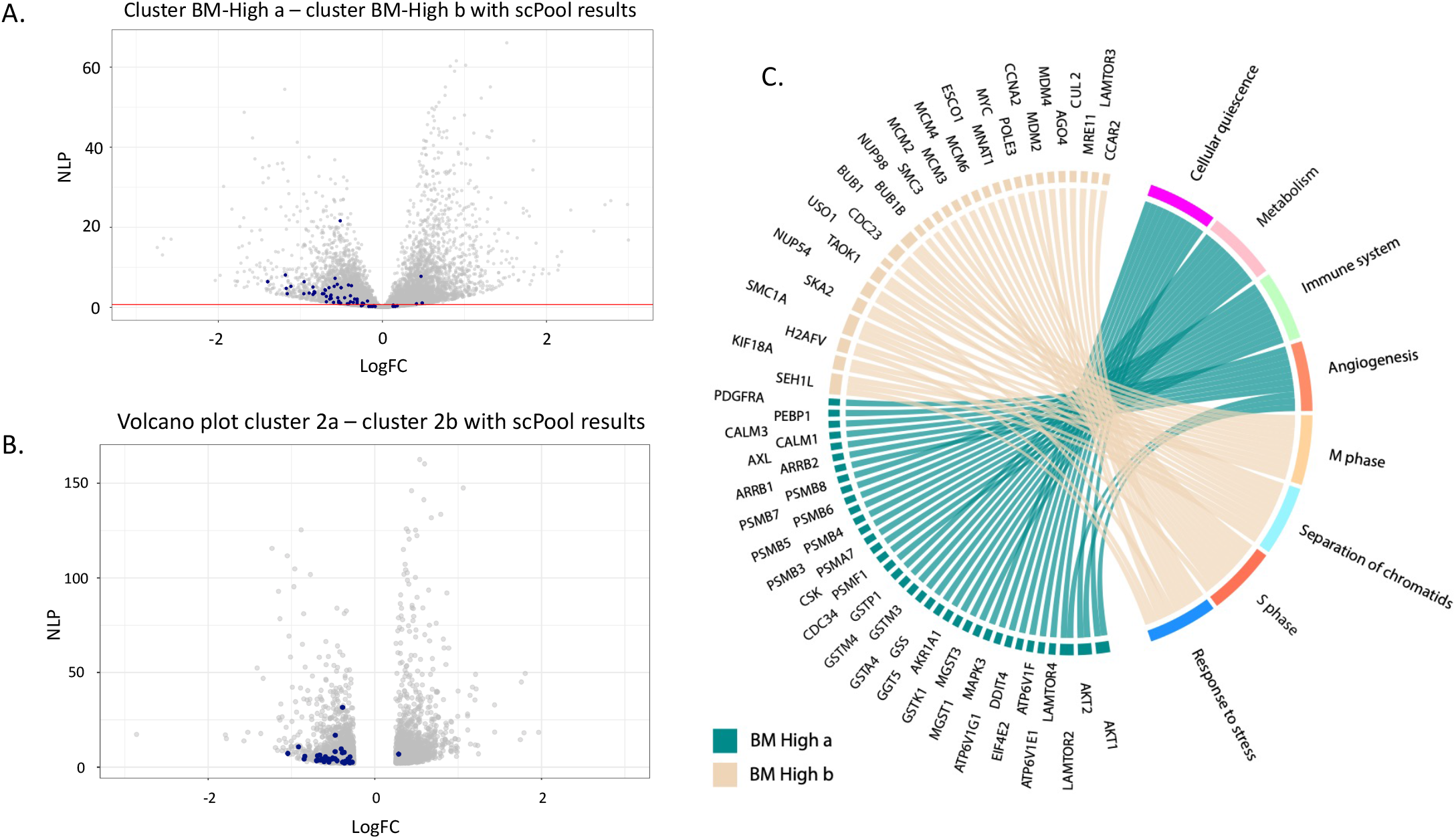
Differential expression between the two High clusters of Bone Marrow-MSC samples. (A) Volcano plot of negative log P-value (NLP) against fold change, created with standard single cell level computation in edgeR. (B) Corresponding volcano plot based on the pseudo pools results. Blue shading indicates cell cycle markers obtained from cycleBase(45). (C) Chord diagram of gene ontology analysis highlighting the top 10 differentially expressed genes in each of 8 pathways representative of the up- and down-regulated genes. Ribbons link genes on the left to pathways on the right; genes associated with multiple pathways bifurcate. Note that in this depiction, the direction of differential expression is the same for all genes in the pathway.

To implement the scPool strategy, we created random pools of pseudo-cells consisting of the sum of raw read counts for random draws of 20 cells (excluding the low-quality cluster 1 cells) within a donor’s sample. The number of such pseudo cells ranged from 6 for sample BM5 to 20 for sample BM4. We then retained all genes with at least one read in 90% of the pseudo cells, a total of 6,422 genes. This dataset was normalized using the supervised normalization of microarrays (SNM) protocol (27) with cluster sub-type as the biological variable, adjusting for donor effects, and analysis of variance was used to detect differentially expressed genes. The procedure was repeated ten times, and the fold change and p-values were averaged to generate a robust list of cluster-specific genes. Compared with the single cell analysis, 1,290 more significant differentially expressed genes were detected. Figure 2C shows generally higher significance than the single cell-level analysis, without over-estimation of fold-changes for a large fraction of the less-significant genes.

Next, we probed the magnitude of donor contributions to variance within clusters, by performing analysis of variance with donor as the fixed effect of interest. Within just the high-quality cluster BM-High_a cells, donor effects accounted for 8.5% of the variance. A similar result was observed for cluster BM-High_b.

### Donor Effects on Umbilical Cord Tissue Derived MSC Gene Expression

Umbilical cord tissue derived MSC (UCT-MSC) samples from four donors were used for scRNA-seq analyses. A total of six scRNA-seq samples were prepared, all from the same lab, including three biological replicates of one donor sample (UCT1a, UCT1b and UCT1c). An average of 349 cells were profiled per sample, with an average read depth of 27,417, representing 9,700 UMI and 3,057 expressed genes per cell (Table 2). The UCT-MSC gene expression profiles were also analyzed with the SC3 pipeline.

Two major clusters of single cell profiles were again observed in the projection of the first two principal components of the UCT-MSC data (Figure 3A). The smallest of these, consisting of 13% of the cells, was characterized by cells with low UMI counts, typically fewer than 2,000 detected genes (Figure 3B), similar to the BM-MCS analysis. These low UMI-count cells were present in every sample but again were more prevalent in two of the samples (UCT1c and UCT3: Figure 3C). It is not clear whether the origin of these cells is a technical artefact, or has a biological basis, but they also appear to be of low quality and were again excluded from all subsequent analyses. Within the high-quality cells, SC3 once more identified two clusters of cells, UCT-High_a (UCT2 and UCT3) and UCT-High_b (UCT1a, UCT1b, UCT1c and UCT4), though in this case they did not clearly correspond to one of the Principal Components. The three replicates of donor UCT1 were primarily captured within the UCT-High_b cluster, suggesting consistency of technique.

**Figure 3.**
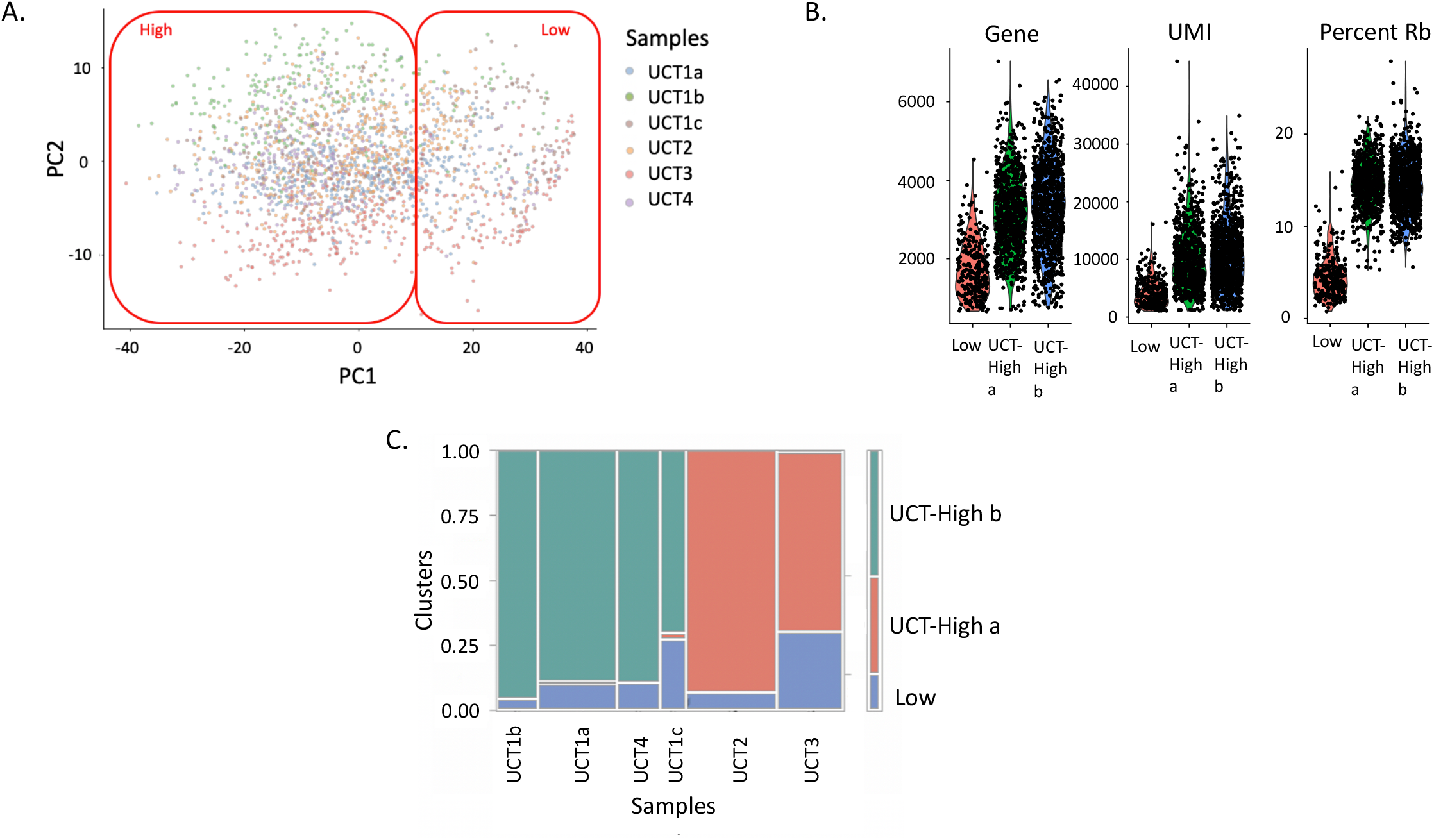
Clusters of UCT-MSC Profiles. (A) The first two Principal Components of gene expression identify the two major clusters of cells, which are colored by sample. Most cells of each sample cluster together. (B) Violin plots show the density of the number of Genes, UMI, and Ribosomal Protein transcripts (RP) per cell. (C) Association of cells with clusters. The width of each column is proportional to the number of cells in the indicated sample, and the color of each box corresponds to cells in cluster 1 (blue), 2a (red) or 2b (green).

Implementation of the scPool strategy for detecting differential gene expression between the two UCT-MSC clusters, after removing the low-quality cells, detected 2,526 genes at an FDR of 5%. Directional up-regulation of established marker genes for mitosis is evident in cluster UCT-High_b as indicated by blue points in the volcano plot Figure 4A. Gene ontology analysis (Figure 4B) indicates enrichment for up-regulation of collagen biosynthesis, integrin signaling, extracellular matrix (ECM) organization, and protein translation pathways in the cluster UCT-High_a MSCs, whereas the cluster UCT-High_b cells are enriched for cell cycle regulation, degradation of mitotic proteins, as well as various processes related to cell cycle progression, including *CDC20* mediated degradation of Securin, and auto degradation of *CDH1*, suggesting potential donor-dependent heterogeneity in the gene expression profile of UCT-MSC.

**Figure 4.**
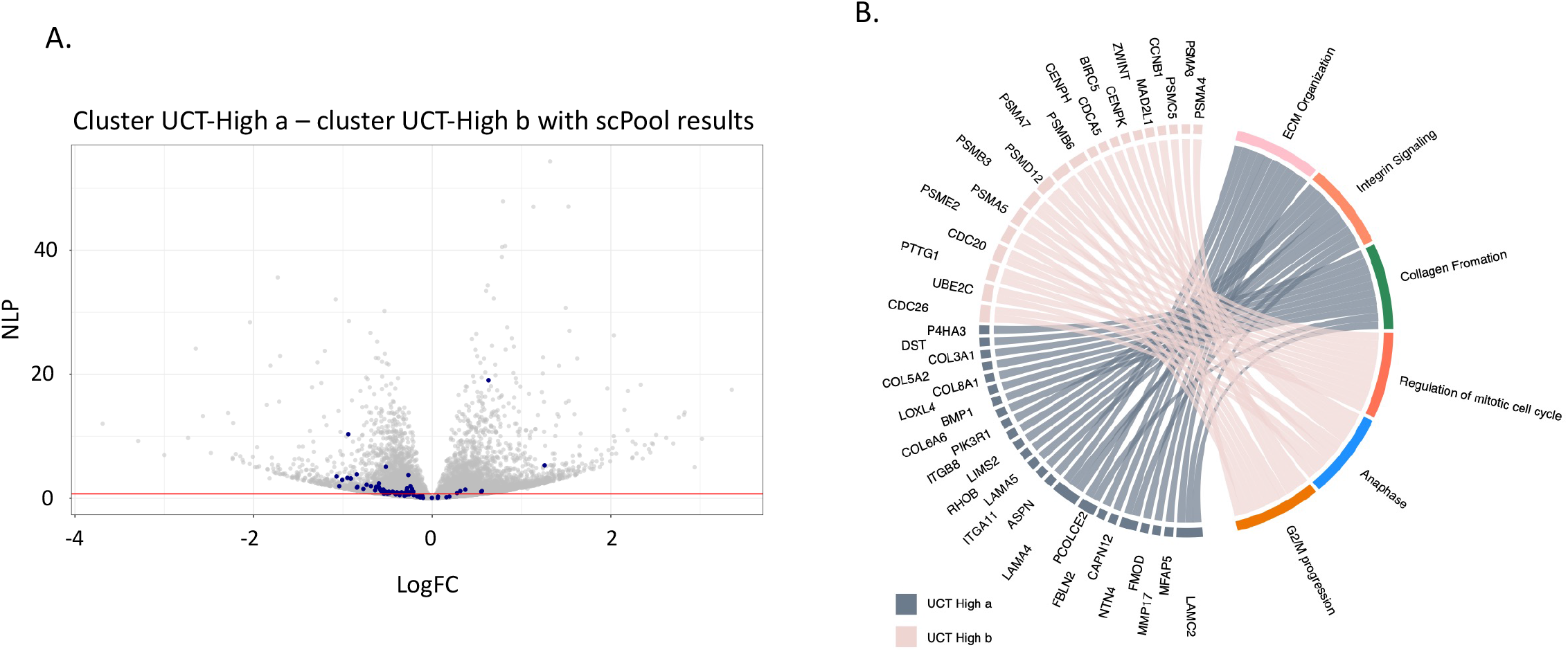
Differential expression between two high quality UCT-MSC clusters. (A) Volcano plot of significance against fold difference in gene expression for the comparison of clusters 2a and 2b. Blue points indicate genes with established roles in cell-cycle regulation. (B) Chord diagram of gene ontology analysis highlighting the top 10 differentially expressed genes in each of 10 pathways representative of the up- and down-regulated genes. Ribbons link genes on the left to pathways on the right; genes associated with multiple pathways bifurcate.

### Comparison between Bone-Marrow and Cord-Tissue derived MSC single-cell gene expression profiles

Direct comparison of MSCs derived from bone marrow and MSCs from umbilical cord tissue was performed by combining the analyses of the previous two datasets. As expected, Principal Component Analysis (PCA) of the raw single cell profiles again identified two major clusters of low and high UMI-abundance cells along PC1, but in this case PC2 of the joint analysis cleanly differentiates the BM and CT samples (Figure 5A). This result implies that there are significant differences in gene profile between MSCs derived from the two source tissues.

**Figure 5.**
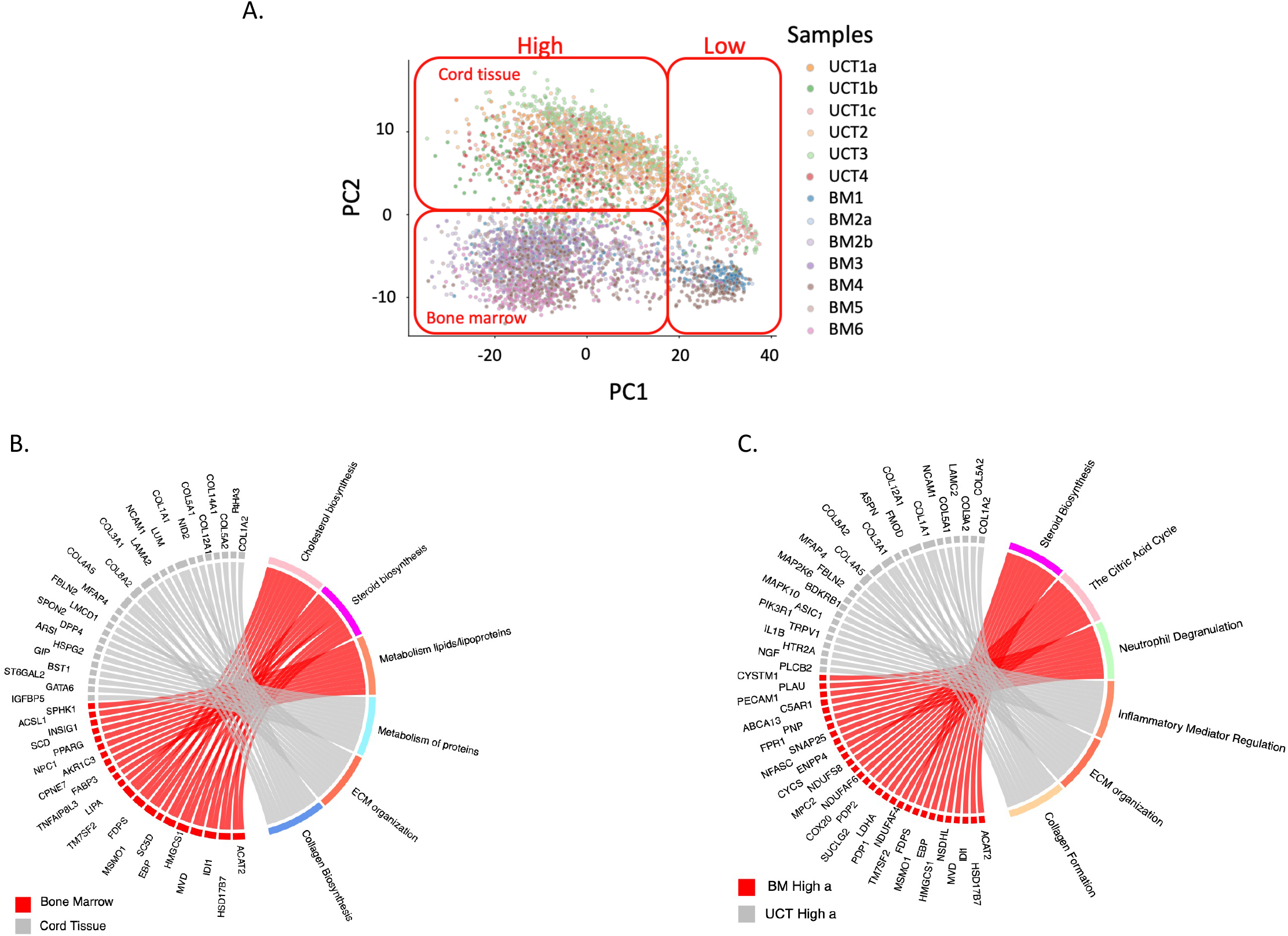
(A) Principal component analysis showing clustering of MSC samples by tissue of origin (bone marrow vs umbilical cord tissue). (B) Chord diagram summarizing differential expression between UCT-MSC and BM-MSC. (C) Cluster-specific pathway expression is not conserved between the two High clusters from the two tissues of MSC origin. Chord diagram contrasting clusters BM-High_a versus UCT-High_a.

We implemented scPool to identify differentially expressed genes between MSCs derived from the two sources, after removing the low-quality cells identified in our first series of analysis. Pools of 20 cells were again assembled computationally within donor and mixed model analysis of variance was performed, with donor as a random effect. 3,437 genes were found to be up-regulated in the BM-MSC, and 3,250 genes down-regulated, compared to UCT-MSCs. We highlighted the top 16 differentially expressed genes (Figure S4). Gene ontology analysis of all the significantly differential expressed genes indicates enrichment for metabolism of lipids and lipoproteins, cholesterol biosynthesis, mitochondrial translation, and metabolic pathways in the BM-MSCs, whereas the UCT-MSCs were enriched for ECM organization, collagen biosynthesis and signal transduction (Figure 5B). UCT-MSC also showed relative up-regulation of mitotic cell cycle pathways, but this likely reflects the greater ratio of UCT-High_b to UCT-High_a cluster cells than of BM-High_b to BM-High_a cells, rather than a consistent trend favoring cell division in the UCT-MSC.

Pathway enrichment analysis also showed that the differences between the two clusters in each tissue are not consistently maintained. The chord diagram (Figure 5C) highlights pathways overexpressed in BM-High_a and UCT-High_a, which are not the same. These data confirm differences in the gene expression of non-dividing cells as a function of tissue of origin and suggest that the two types of MSCs are likely to have divergent regulatory and functional potentials. Importantly, the UCT-High_a population exhibit higher expression of genes involving pro inflammatory mediation, ECM organization and collagen biosynthesis, whereas the BM-High_a population had higher expression of steroid biosynthesis, the citric acid cycle and neutrophil degranulation genes (Figure 5C).

Next, we examined the expression level of genes that play important roles in the immunomodulatory response induced by MSCs. Focused comparison of expression of genes that are associated with cell adhesion, migration, immunosuppression and immunostimulation between the BM- and UCT-derived MSCs suggests tissue-of-origin and donor differences in gene activity (Figure S5). BM-High_a cells characteristically overexpress transcripts encoding the membrane proteins prostaglandin synthase (*PTGES2*) and Endoglin (*ENG*) as well as the lysosomal protein CD63, compare to BM-High_b, whereas BM-High_b cells overexpress the genes *CD46* (encoding a complement cofactor), *CD47* (an integrin-associated protein), and *CD146* (*MCAM*, cell adhesion molecule), compare to BM-High_a. These genes are in general overexpressed in BM derived MSC compare to UCT derived MSC. BM derived MSCs have higher expression of the cell surface glycoprotein coding genes *CD44* and *CD59*, as well as the nucleotidase *NT5E* and immune checkpoint molecule *CD276* compare to UCT derived MSC. Conversely, when comparing UCT-High_a and UCT-High_b clusters, UCT-High_a cells overexpress the tetraspanin regulators of motility *CD151*, and cell surface protein coding genes *CD99, THY1* and *CD248*, while UCT-High_b overexpressed *CD9*. The cell surface protein coding gene *CD81* is not significantly differentially expressed between the clusters UCT-High_a and UCT-High_b. We looked at two genes associated with immunostimlation: *CCL2* and *CD109*, which are overexpressed in UCT and BM derived MSCs, respectively.

We also looked at multiple pluripotent and stemness genes (Figure S6), none of which were found to be significantly differentially expressed between bone marrow and umbilical cord tissue.

## Discussion

MSCs from various tissue sources are the subject of 4044 registered clinical trials (ClinicalTrials.gov-keyword: MSC as other terms with filters - not yet recruiting, recruiting, enrolling by invitations, and active, not recruiting; search date – August 7, 2020). It is thus important to develop robust high-throughput approaches for characterization of diverse batches from various tissue sources in order to help evaluate reasons for success or failure of individual trials or patient responses. Single cell RNA sequencing is a relatively unbiased approach to profile the molecular attributes of individual cells. Potential utility of scRNA-seq includes characterization of heterogeneity that cannot be observed with bulk RNA-seq, and monitoring of the effect of the stage of the cell-cycle on transcriptional diversity.

Contrasting pre-freeze and post-thaw samples from 2 donors, we identified numerous differentially expressed genes that are associated with different types of cellular functions, such as cytokine signaling, cell proliferation, cell adhesion, cholesterol/steroid biosynthesis, and regulation of apoptosis. Previuosly, functional differences between pre-freeze (fresh) and post-thaw MSCs were also reported by others (28). In this study, however, we focused on in-depth scRNA-seq analysis of post-thaw MSCs as they are currently being tested as cell therapy products in many clinical trials.

Here we describe a droplet-based scRNA-seq comparison of donor, tissue-of-origin, and expansion conditions of out-of-thaw MSC variability, concluding that bone marrow and umbilical cord tissue-derived MSCs have significant differential expression that likely explains some of the documented differences between them, and that donor differences are modest yet significant. To our knowledge, six other scRNA-seq studies (11, 12, 20-23) of MSCs have been published, and our results are broadly concordant though with some important differences in emphasis. Barrett *et. al*., 2019 (22) used a version of SmartSeq to deeply profile 103 Wharton’s jelly-derived umbilical cord MSCs and 63 bone marrow-derived MSCs, identifying 463 differentially expressed genes enriched for activity in numerous processes including the matrisome, coagulation, angiogenesis, and wound-healing via immune-regulation. The current study similarly finds a difference between cord tissue and bone marrow-derived MSCs. Additionally, our data also shows a cell cycle variability which seems to be related to the donor (22).

Each of the other studies (11, 12, 20-23) has noted that the cell cycle gene expression is a major source of heterogeneity within donors. According to the *Huang et. al*., 2019 (20) study, it appears the cell cycle is related to the immune regulatory potency of the MSCs. Previous work (21) has used a core set of G1/G2M/S markers to assign cells to each phase, and regressed out this source of variance before performing downstream analysis. We eschewed this approach both because of concerns over the reliability of the assignments, and to emphasize that the proportion of cells with low expression of these genes is an important component of among-donor differences in both BM-MSC and UCT-MSC. Reported higher proliferative capacity of Wharton’s jelly-derived MSCs (22) is consistent with the higher proportion of mitotic genes in our UCT-MSCs relative to BM-MSCs. However, it should be emphasized that higher overall expression may not correlate with higher rates of proliferation, since expression levels may vary among donors without implying that a different fraction of cells are undergoing division. On the other hand, it appears that putative G0 cells that do not express cell cycle genes have quite different transcriptional properties that are directly relevant to their biological functions such as immunomodulatory potential. We note that each of our samples was profiled at population doubling level (PDL) ranged from 12-15, eliminating passage number as a source of variability in our study.

Other authors have also chosen to regress out “batch” effects before searching for heterogeneity, even though in each case “batch” appears to be coincident with “donor” (8, 21) or “Passage” (20). In the absence of biological replication, that is, two MSC preparations obtained independently from the same donor, it is impossible to know whether differences between sample populations have a biological or technical basis. Nevertheless, we estimate from principal component variance analysis that less than 10% of the overall expression variability is among donors/batches within each of the two clusters observed in both the BM- and UCT-MSC datasets (Figure 1C, Figure 3C). We see this minor source of variability is donor-related in the two instances where we had technical replicates from the same donor (in the case of the two BM-MSC samples cultured in different laboratories). The cells strongly tended to be assigned to the same sub-cluster BM-High_a or UCT-High_a. Whether or not these differences impact MSC function in clinical applications remains to be seen, additional large-scale comparisons with a large set of samples with high quality data on patient outcomes will need to be analyzed.

In this study the transcriptomes of human bone marrow and cord tissue-derived MSCs were analyzed via drop-seq single cell RNA-seq. Using this approach, new information about MSCs emerges. First, the differences between bone marrow-derived MSCs and cord-tissue derived MSCs were seen. Surprisingly, pathways up-regulated in G0 bone marrow-derived MSCs did not correspond to the same pathways upregulated in G0 cord tissue -derived MSC (Figure 5C). Further, we observed differences in various immune regulatory genes between bone marrow and cord tissue MSCs, especially for the “a” cluster cells (Figure 5C). Notably, BM-High_a MSCs had higher gene expression for *PTGES2*, and the protein encoded by this gene is known to have direct or indirect role in immunomodulation by MSCs (29, 30). *PTGES2* encodes membrane-bound prostaglandin synthase E2 which converts prostaglandin H2 (*PGH2*) to prostaglandin E2 (*PGE2*) that is known to have anti-inflammatory/immunosuppressive effects on various immune cells, including macrophages, T cells and B cells (31, 32).

MSC surface proteins are important for their significant roles in identification and functions (33). When we compared gene expression for surface markers that are known to have some immunomodulatory functions, BM derived MSCs showed higher expression for *CD46, CD47, and CD276* whereas UCT derived MSC had higher expression for *CD81*. Surface expression of CD46 protein helps MSCs to inhibit complement binding and complement-mediated lysis (34). *CD47* serves as a “don’t eat me” signal to avoid phagocytosis by engaging its cognate ligand signal-regulatory-protein alpha (SIRP alpha) on phagocytes (35, 36), and the interaction of *CD47* with SIRP alpha is reported to inhibit antigen presenting cell (APC) maturation and enhance STAT3 phosphorylation and IL10 induction in APC (37). *CD276* is known to cause immune suppression by inhibiting T cell function and is currently being targeted as a check point blockade therapy for cancer (38); however, their specific role in MSC-mediated immunomodulation is not yet confirmed. CD81 is one of the surface markers used to identify MSC-derived extracellular vehicles (EVs) but does not have any reported immunomodulatory role for MSCs; however, *CD81* coding gene is known to affect T regulatory (Treg) and myeloid-derived suppressor cell (MDSC) function enhancing tumor growth (39). Taken together, gene expression differences for surface markers related to immune response between BM and UCT-MSCs implicates potential differences in the immunomodulatory functions between BM and UCT-MSCs. Further, differences in immunomodulatory gene expression between the High_a and High_b clusters for both BM and UCT-MSCs indicates functional and phenotypic heterogeneity within each MSC product. No differences in expression of a small number of pluripotent and stemness marker genes was detected between BM and UCT derived MSCs (Figure S6), though we note that abundance of these transcripts was very low which reduces power to observe differential expression.

In summary, this study both confirms the potential for functional differences to exist between MSCs derived from different tissues and even donors, and that within-sample heterogeneity is low. The expression of cell cycle markers is a major component of heterogeneity among donors, and manufacturing processes may need to accommodate biological and technical influences on proliferative potential. These findings will help improve the therapeutic MSC manufacturing processes and identify the most efficient cells from a heterogeneous MSC population. Even though differences in the gene expression profile between bone marrow and cord tissue G0 MSC were found, further studies are needed to confirm these results as well as the impact of these differences on the clinical use of these cells.

## Experimental Procedures

### Study approval

This study was approved by the ethics committee of the institutional review boards at Georgia Institute of Technology and Duke University (IRB protocol no. H17348). All procedures involving human participants were in accordance with the ethical standards of the research committee. Informed consent was obtained from all participants.

### Human Bone Marrow MSC collection

Seven frozen human bone marrow-derived MSC lots from six male donors were purchased from RoosterBio Inc., and expanded using RoosterBio expansion protocol (https://www.roosterbio.com/wp-content/uploads/2019/10/A.-RoosterBio-MSC-001-BOM-Expansion-Protocol-IF-08022016.pdf). Briefly, a BM-hMSC high performance media kit was brought to room temperature. Then 1 vial of Media Booster GTX (RoosterBio, catalog no. SU-003) was added to 500ml hMSC high performance basal media (RoosterBio, catalog no. SU-005). The 10million BM-hMSC vial was obtained from a liquid nitrogen dewar and immediately thawed at 37°C for approximate 2 minutes while monitoring the process and removed from the water bath once a small bit of ice remained. Cells were aseptically transferred to a 15ml centrifuge tube and 10ml cultured media was added. The cells were spun down at 200g for 10 minutes and all the supernatant was discarded. The cell pellet was re-suspended in 10ml of culture media and transferred into 500ml media bottle. The cells were mixed by capping and gently inverting the bottle and distributed (seeded at 3500-4000 cells/cm^2^ and 42 mL media/T225) equally into T-225 vessels (Corning cat no. 431082). The vessels were transferred to a 37°C incubator ensuring that the surfaces were covered with media. The vessels were observed microscopically from day 1 to determine percentage confluency. Once they reached >80% confluency, they were harvested the next day, and cryo-preserved in Cryostor CS-10 freezing media. All single-cell RNA-sequencing was performed on the out of thaw cells directly from these frozen vials.

Samples BM1 and BM2a were cultured in Laboratory A, while samples BM2b, BM3, BM4, BM5 and BM6 were cultured in Laboratory B. Samples BM2a and BM2b were from the same donor.

### Human Cord Tissue MSC collection

For cord tissue derived samples, six frozen human MSC samples from four male donors were provided by the department of pediatrics, Duke University. Cryopreserved P0 vials were placed in a sterile bag which was itself placed in a 37°C water bath. Vials were thawed until the cell suspension was slushy (∼2 minutes). Cell suspension from the vials were transferred to a 15 mL tube containing XSFM (Irvine Scientific, cat. no. 91149) using a sterile serological pipette and the cell suspension was mixed slowly. The cryovial was rinsed with 0.5 mL of XSFM and the rinse was transferred to the 15 mL tube. After mixing slowly, the cell count and viability was measured. The cells were mixed in the 15 mL conical using a sterile pipette and transfer the volume containing 3.4 × 10^6^ cells into a HYPER flask containing 1.12 L of XSFM and the bottle was mixed gently. The HYPER Flasks were placed into a 37°C/5% CO2 incubator. The P1 cells were harvested after 5-7 days. The P2 cells were then frozen using CS-10 freezing media and cryopreserved. The Cryopreserved P2 cells were shipped to us for the downstream characterizations. All the single-cell RNA-sequencing was performed on the out of thaw cells directly from these frozen vials.

Samples UCT1a, UCT1b and UCT1c come from the same donor.

### Thawing and single cell suspension preparation for Single-Cell RNA-Sequencing

Frozen vials containing 1 million MSC were thawed in a 37°C water bath for a couple of minutes. Cells were then aseptically transferred to a 15 ml centrifuge tube. Room temperature RPMI media (1mL) was used to rinse the cell vial and added to the cells in the 15 ml tube. Another 3 ml of media was added to the cells and mixed well with serological pipette. Cells were counted using Nucleocounter and spun down at 200 g for 10 minutes. The cells were re-suspended in media and counted again and processed for scRNA-sequencing.

### Single-cell RNA-seq library preparation and sequencing

The Illumina-Bio-Rad ddSEQ platform was used to process, capture, and barcode the cells to generate single-cell Gel Beads by following the manufacturer’s protocol. Cell suspensions were loaded onto a ddSEQ Cartridge along with reverse transcription master mix, and encapsulated and barcoded by the Single-Cell Isolator. Lysis and barcoding took place in each droplet. Droplets were disrupted and cDNA was pooled for second strand synthesis. Libraries were generated with direct cDNA tagmentation using Nextera technology. Tagmentation was followed by 3′ enrichment and sample indexing to prepare indexed, sequencing-ready libraries. The libraries were sequenced using Nextseq sequencing Platform on an Ilumina NextSeq in the IBB Molecular Evolution core at Georgia Tech (PE75, mid-output V2.5 kit). The library quality (check for primer dimers, adopter dimers, ethanol contamination, degradation as well as the size and concentration) was confirmed before each sequencing run using Agilent Bioanalyzer2100.

All the BM-MSC samples have 2415 cells with an average 3,835 genes/cell (Table 1) and the UCT-MSCs have 1785 cells with average 3056 genes/cell (Table 2). The BM-MSC samples have average 28,890,505 (Table 1) reads per sample and the UCT-MSC samples have 33,771,805 (Table 2) reads per sample in average.

Confirmation that MSCs were relatively pure populations of undifferentiated cells was revealed by FACS analysis of cell surface markers provided by the manufacturer as a release criteria. Furthermore, scRNA-seq (which is less sensitive due to high drop-out rates) confirmed prevalent expression of *NT5E* (CD73), *THY1* (CD90), and *ENG* (CD105) and absent expression of *CD34* among other genes (Fig. 6). *ENG* was expressed on 66% of the cell, *NT5E* on 72%, and *THY1* on 92%. In contrast, each of the transcripts *PTPRC, CD34, CD14, ITGAM, CD79A, CD19*, and *HLA-DRA* were detected in less than 0.5% of the cells.

### Data Analysis

Sample demultiplexing and gene counts were extracted using the Illumina Sure cell pipeline. The raw reads were trimmed, and the gene-barcode matrix was generated. Sure cell was also used to filter and align the samples and to generate gene-cell UMI count matrices. The seven samples from bone marrow were sequenced in three different batches, and the six samples from cord tissue were sequenced in two batches.

Downstream analysis was initiated with SC3 software (40) for cell clustering. Owing to the high proportion of zero counts in most cells (as is typical of dropseq data), we elected to perform differential expression analysis on pools of pseudo-cells whose profiles have a distribution of read counts that closely resembles that of bulk RNAseq. Custom R scripts were used to generate pools of pseudo-cells by pooling groups of 20 cells within each sample and cluster, and then summing their gene count. The gene expression values from the pseudo cells were normalized to counts per million before using EdgeR (41) for normalization and differential expression estimation. Default likelihood ratio tests assuming negative binomial distributions were performed in EdgeR to evaluate the significance of differential expression. Ten permutations of this procedure were performed, and the average differential expression and negative-log P-values were computed, and genes significant with a FDR less than 5% were selected for downstream gene ontology analysis. Then gene ontology was performed on the differentially expressed genes using GSEA (42, 43) and ToppGene (44) tools.

## Acknowledgements

This work was funded by the Billie and Bernie Marcus Foundation with grants to the Marcus Center for Therapeutic Cell Characterization and Manufacturing (MC3M) at Georgia Tech and the Marcus Center for Cellular Cures (MC3) at Duke University. We thank Georgia Research Alliance for their support for this study. All the samples were sequenced on Illumina Nextseq500 platform at Georgia Institute of Technology Molecular Evolution Core to generate single cell RNAseq data. KR is partially funded by the NSF Engineering Center for Cell Manufacturing Technologies (CMaT) though grant EEC 1648035.

## Author contributions

JK contributed on the UCT-MSC culture. MEO expanded and cryopreserved some of the BM samples under the supervision of EAB. PC generated the scRNA-seq data, which was primarily analyzed by CMT under the supervision of GG. PP curated the pathway analyses and compared with prior studies. CY and KR conceived the experiments and supervised all aspects of the study. CMT, PC and GG wrote the first draft, and all authors revised the paper.

## Declaration of Interest

The authors have declared that no conflict of interest exists.

**This article has not been yet accepted or published anywhere else**.

## References

1. Brown C, McKee C, Bakshi S, Walker K, Hakman E, Halassy S, et al. Mesenchymal stem cells: Cell therapy and regeneration potential. J Tissue Eng Regen Med. 2019;13(9):1738–1755.

2. Galipeau J, and Sensebe L. Mesenchymal Stromal Cells: Clinical Challenges and Therapeutic Opportunities. Cell Stem Cell. 2018;22(6):824–833.

3. Loebel C, and Burdick JA. Engineering Stem and Stromal Cell Therapies for Musculoskeletal Tissue Repair. Cell Stem Cell. 2018;22(3):325–339.

4. Kim H, Bae C, Kook YM, Koh WG, Lee K, and Park MH. Mesenchymal stem cell 3D encapsulation technologies for biomimetic microenvironment in tissue regeneration. Stem Cell Res Ther. 2019;10(1):51.

5. Richardson SM, Kalamegam G, Pushparaj PN, Matta C, Memic A, Khademhosseini A, et al. Mesenchymal stem cells in regenerative medicine: Focus on articular cartilage and intervertebral disc regeneration. Methods. 2016;99:69–80.

6. Mastrolia I, Foppiani EM, Murgia A, Candini O, Samarelli AV, Grisendi G, et al. Challenges in Clinical Development of Mesenchymal Stromal/Stem Cells: Concise Review. Stem Cells Transl Med. 2019;8(11):1135–1148.

7. Dominici M, Le Blanc K, Mueller I, Slaper-Cortenbach I, Marini F, Krause D, et al. Minimal criteria for defining multipotent mesenchymal stromal cells. The International Society for Cellular Therapy position statement. Cytotherapy. 2006;8(4):315–317.

8. Liu X, Xiang Q, Xu F, Huang J, Yu N, Zhang Q, et al. Single-cell RNA-seq of cultured human adipose-derived mesenchymal stem cells. Sci Data. 2019;6:190031.

9. Lo Surdo JL, Millis BA, and Bauer SR. Automated microscopy as a quantitative method to measure differences in adipogenic differentiation in preparations of human mesenchymal stromal cells. Cytotherapy. 2013;15(12):1527–1540.

10. Samsonraj RM, Rai B, Sathiyanathan P, Puan KJ, Rotzschke O, Hui JH, et al. Establishing criteria for human mesenchymal stem cell potency. Stem Cells. 2015;33(6):1878–1891.

11. Macosko EZ, Basu A, Satija R, Nemesh J, Shekhar K, Goldman M, et al. Highly Parallel Genome-wide Expression Profiling of Individual Cells Using Nanoliter Droplets. Cell. 2015;161(5):1202–1214.

12. Zheng GX, Terry JM, Belgrader P, Ryvkin P, Bent ZW, Wilson R, et al. Massively parallel digital transcriptional profiling of single cells. Nat Commun. 2017;8:14049.

13. Phinney DG. Biochemical heterogeneity of mesenchymal stem cell populations: clues to their therapeutic efficacy. Cell Cycle. 2007;6(23):2884–2889.

14. Buenrostro JD, Corces MR, Lareau CA, Wu B, Schep AN, Aryee MJ, et al. Integrated Single-Cell Analysis Maps the Continuous Regulatory Landscape of Human Hematopoietic Differentiation. Cell. 2018;173(6):1535–1548 e1516.

15. Psaila B, Barkas N, Iskander D, Roy A, Anderson S, Ashley N, et al. Single-cell profiling of human megakaryocyte-erythroid progenitors identifies distinct megakaryocyte and erythroid differentiation pathways. Genome Biol. 2016;17:83.

16. Velten L, Haas SF, Raffel S, Blaszkiewicz S, Islam S, Hennig BP, et al. Human haematopoietic stem cell lineage commitment is a continuous process. Nat Cell Biol. 2017;19(4):271–281.

17. Papalexi E, and Satija R. Single-cell RNA sequencing to explore immune cell heterogeneity. Nat Rev Immunol. 2018;18(1):35–45.

18. Villani AC, Satija R, Reynolds G, Sarkizova S, Shekhar K, Fletcher J, et al. Single-cell RNA-seq reveals new types of human blood dendritic cells, monocytes, and progenitors. Science. 2017;356(6335).

19. Bjorklund AK, Forkel M, Picelli S, Konya V, Theorell J, Friberg D, et al. The heterogeneity of human CD127(+) innate lymphoid cells revealed by single-cell RNA sequencing. Nat Immunol. 2016;17(4):451–460.

20. Huang Y, Li Q, Zhang K, Hu M, Wang Y, Du L, et al. Single cell transcriptomic analysis of human mesenchymal stem cells reveals limited heterogeneity. Cell Death Dis. 2019;10(5):368.

21. Sun C, Wang L, Wang H, Huang T, and Zhang X. Single-cell RNA-seq highlights heterogeneity in human primary Wharton’s Jelly mesenchymal stem/stromal cells cultured <em>in vitro</em>. bioRxiv. 2019:723130.

22. Barrett AN, Fong CY, Subramanian A, Liu W, Feng Y, Choolani M, et al. Human Wharton’s Jelly Mesenchymal Stem Cells Show Unique Gene Expression Compared with Bone Marrow Mesenchymal Stem Cells Using Single-Cell RNA-Sequencing. Stem Cells Dev. 2019;28(3):196–211.

23. Khong SML, Lee M, Kosaric N, Khong DM, Dong Y, Hopfner U, et al. Single-Cell Transcriptomics of Human Mesenchymal Stem Cells Reveal Age-Related Cellular Subpopulation Depletion and Impaired Regenerative Function. Stem Cells. 2019;37(2):240–246.

24. Moll G, Geissler S, Catar R, Ignatowicz L, Hoogduijn MJ, Strunk D, et al. Cryopreserved or Fresh Mesenchymal Stromal Cells: Only a Matter of Taste or Key to Unleash the Full Clinical Potential of MSC Therapy? Adv Exp Med Biol. 2016;951:77–98.

25. Klein AM, Mazutis L, Akartuna I, Tallapragada N, Veres A, Li V, et al. Droplet barcoding for single-cell transcriptomics applied to embryonic stem cells. Cell. 2015;161(5):1187–1201.

26. Stuart T, Butler A, Hoffman P, Hafemeister C, Papalexi E, Mauck WM, 3rd, et al. Comprehensive Integration of Single-Cell Data. Cell. 2019;177(7):1888–1902 e1821.

27. Mecham BH, Nelson PS, and Storey JD. Supervised normalization of microarrays. Bioinformatics. 2010;26(10):1308–1315.

28. Chinnadurai R, Copland IB, Garcia MA, Petersen CT, Lewis CN, Waller EK, et al. Cryopreserved Mesenchymal Stromal Cells Are Susceptible to T-Cell Mediated Apoptosis Which Is Partly Rescued by IFNgamma Licensing. Stem Cells. 2016;34(9):2429–2442.

29. Regmi S, Pathak S, Kim JO, Yong CS, and Jeong JH. Mesenchymal stem cell therapy for the treatment of inflammatory diseases: Challenges, opportunities, and future perspectives. Eur J Cell Biol. 2019;98(5-8):151041.

30. Ghannam S, Bouffi C, Djouad F, Jorgensen C, and Noel D. Immunosuppression by mesenchymal stem cells: mechanisms and clinical applications. Stem Cell Res Ther. 2010;1(1):2.

31. Harris SG, Padilla J, Koumas L, Ray D, and Phipps RP. Prostaglandins as modulators of immunity. Trends Immunol. 2002;23(3):144–150.

32. Aggarwal S, and Pittenger MF. Human mesenchymal stem cells modulate allogeneic immune cell responses. Blood. 2005;105(4):1815–1822.

33. Niehage C, Steenblock C, Pursche T, Bornhauser M, Corbeil D, and Hoflack B. The cell surface proteome of human mesenchymal stromal cells. PLoS One. 2011;6(5):e20399.

34. Le Blanc K, and Mougiakakos D. Multipotent mesenchymal stromal cells and the innate immune system. Nat Rev Immunol. 2012;12(5):383–396.

35. Oldenborg PA, Gresham HD, and Lindberg FP. CD47-signal regulatory protein alpha (SIRPalpha) regulates Fcgamma and complement receptor-mediated phagocytosis. J Exp Med. 2001;193(7):855–862.

36. Jaiswal S, Jamieson CH, Pang WW, Park CY, Chao MP, Majeti R, et al. CD47 is upregulated on circulating hematopoietic stem cells and leukemia cells to avoid phagocytosis. Cell. 2009;138(2):271–285.

37. Toledano N, Gur-Wahnon D, Ben-Yehuda A, and Rachmilewitz J. Novel CD47: SIRPalpha dependent mechanism for the activation of STAT3 in antigen-presenting cell. PLoS One. 2013;8(9):e75595.

38. Picarda E, Ohaegbulam KC, and Zang X. Molecular Pathways: Targeting B7-H3 (CD276) for Human Cancer Immunotherapy. Clin Cancer Res. 2016;22(14):3425–3431.

39. Vences-Catalan F, Rajapaksa R, Srivastava MK, Marabelle A, Kuo CC, Levy R, et al. Tetraspanin CD81 promotes tumor growth and metastasis by modulating the functions of T regulatory and myeloid-derived suppressor cells. Cancer Res. 2015;75(21):4517–4526.

40. Kiselev VY, Kirschner K, Schaub MT, Andrews T, Yiu A, Chandra T, et al. SC3: consensus clustering of single-cell RNA-seq data. Nat Methods. 2017;14(5):483–486.

41. Robinson MD, McCarthy DJ, and Smyth GK. edgeR: a Bioconductor package for differential expression analysis of digital gene expression data. Bioinformatics. 2010;26(1):139–140.

42. Subramanian A, Tamayo P, Mootha VK, Mukherjee S, Ebert BL, Gillette MA, et al. Gene set enrichment analysis: a knowledge-based approach for interpreting genome-wide expression profiles. Proc Natl Acad Sci U S A. 2005;102(43):15545–15550.

43. Mootha VK, Lindgren CM, Eriksson KF, Subramanian A, Sihag S, Lehar J, et al. PGC-1alpha-responsive genes involved in oxidative phosphorylation are coordinately downregulated in human diabetes. Nat Genet. 2003;34(3):267–273.

44. Chen J, Bardes EE, Aronow BJ, and Jegga AG. ToppGene Suite for gene list enrichment analysis and candidate gene prioritization. Nucleic Acids Res. 2009;37(Web Server issue):W305–311.

45. Santos A, Wernersson R, and Jensen LJ. Cyclebase 3.0: a multi-organism database on cell-cycle regulation and phenotypes. Nucleic Acids Res. 2015;43(Database issue):D1140–1144.

