## Supplemental figure S1, S2, S3, S4, S5, S6 for "Single-Cell RNAseq of Out-of-Thaw Mesenchymal Stromal Cells Shows Striking Tissue-of-Origin Differences and Inter-donor Cell-Cycle Variations"

### SUPPLEMENTARY INFORMATION

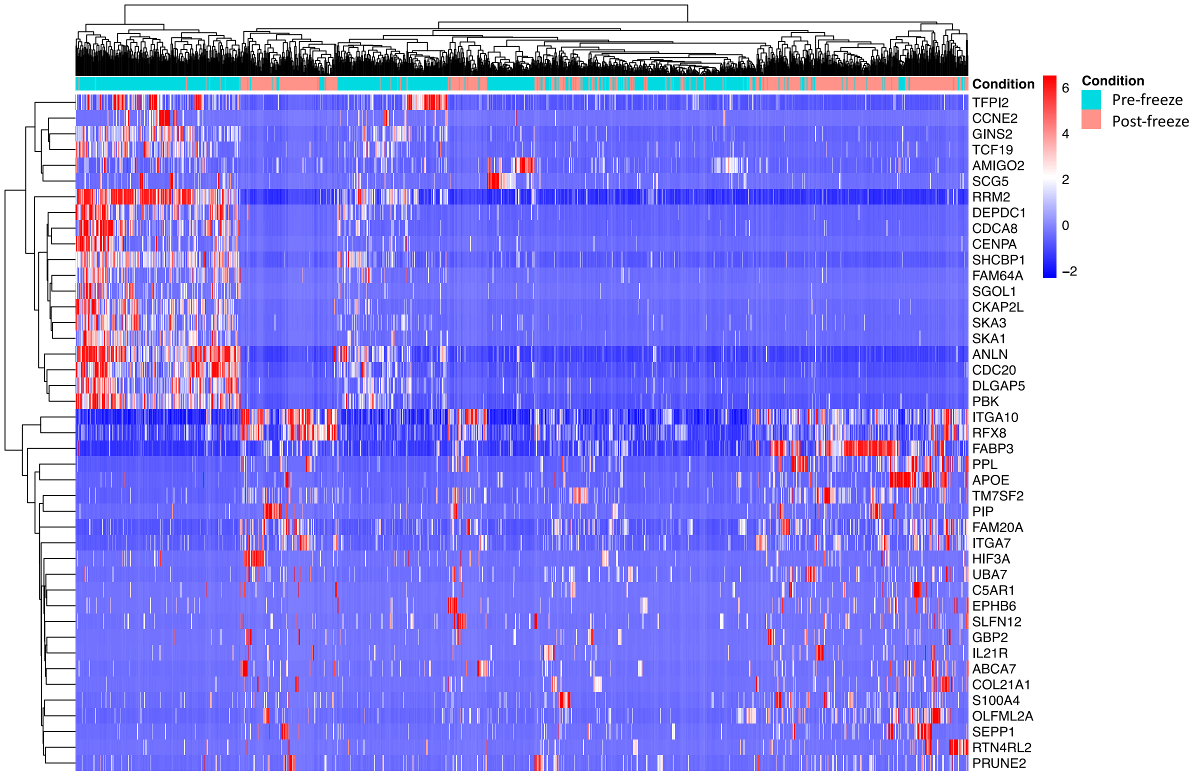


Figure S1: Heatmap displaying the top 50 DE genes for the between post-thaw and pre-freeze MSC comparison. The DEG analysis between these samples shows a significant overexpression of 1,743 genes on the pre-freeze samples, compared to 310 genes significantly overexpressed in the post-thaw samples. The most significant overexpressed pathways on the Pre-freeze samples are cell proliferation and cell adhesion, while the pathways over-expressed in the frozen samples are cholesterol/Steroid biosynthesis and cell death regulation.

A.

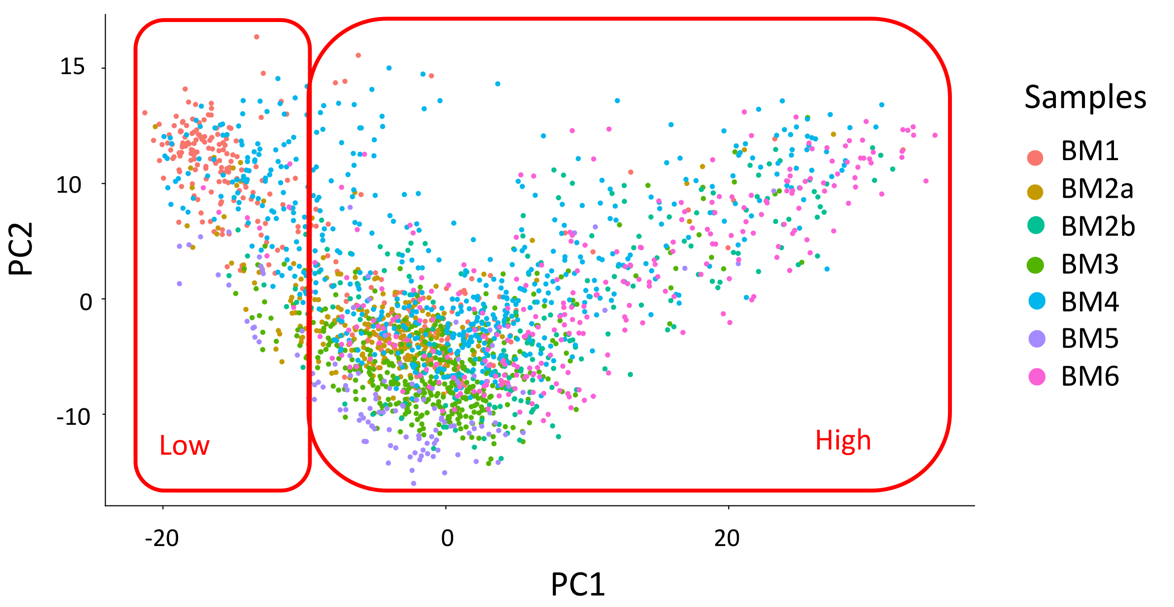


B.

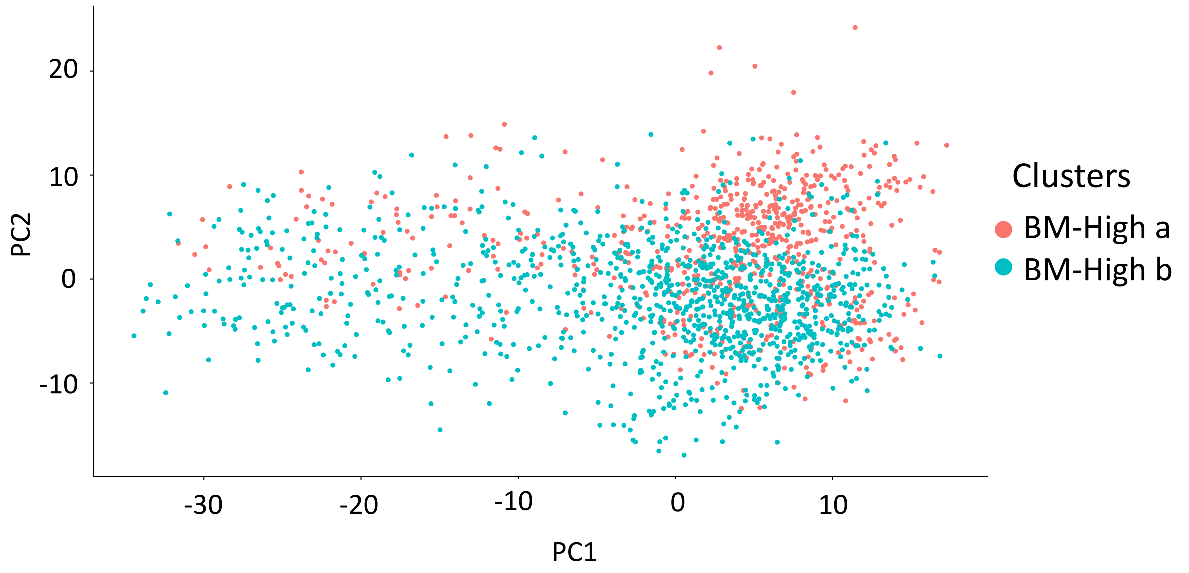


Figure S2: PCAs created with Seurat (A) Two major clusters differentiate along PC1. These clusters correspond to the low (left) and high (right) UMI count cells. (B) This PCA shows the clusters BM-High_a and BM-High_b. This cluster are not as well differentiated when using Seurat as they are when clustering the cells with SC3.

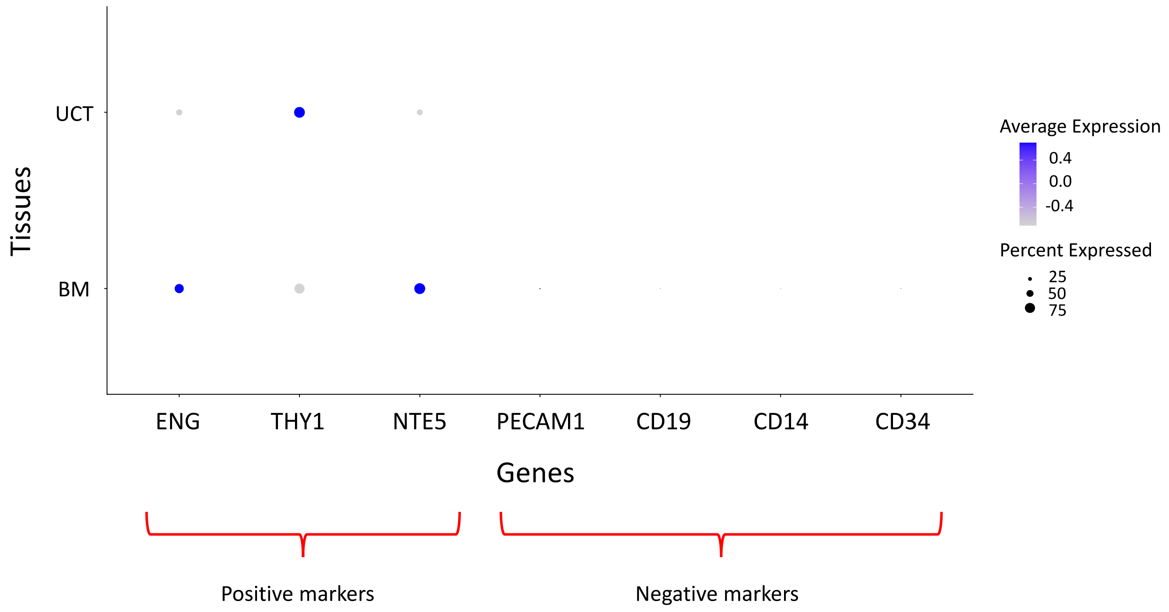


Figure S3: Dot plot displaying the MSC identity markers established by the ISCT. The size of the dots corresponds to the percentage of cells, in the tissue, expressing the gene. The color corresponds to the average non-zero expression of the gene, per cell, in each tissue. Light purple represents low expression per cell, while dark purple corresponds to high expression per cell.

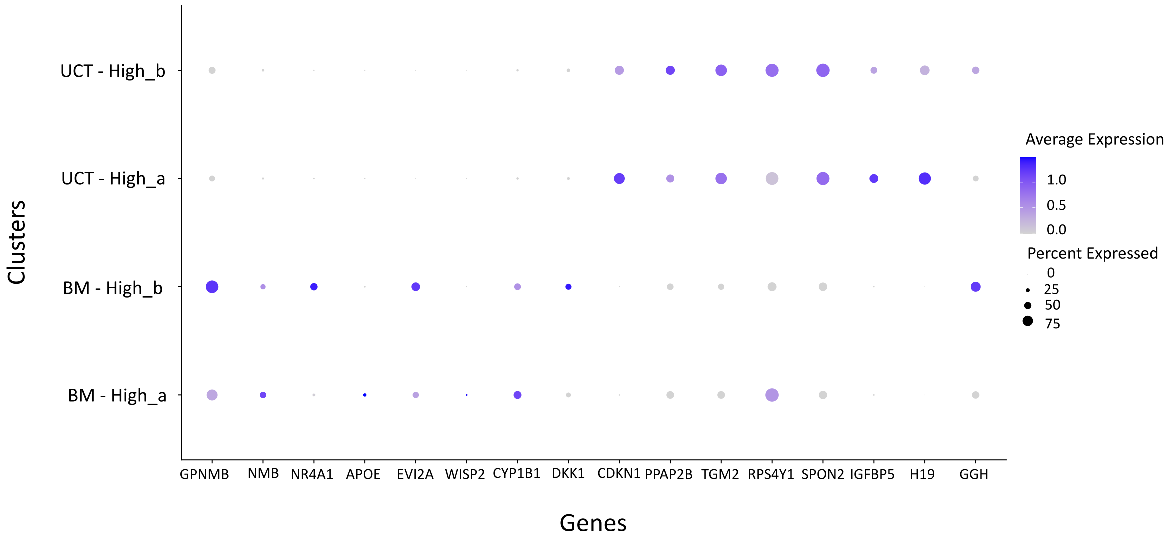


Figure S4: Dot plots displaying the top 16 differentially expressed genes in each group. The first 8 genes are overexpressed in BM derived MSC while the last 8 genes are overexpressed in UCT derived MSCs.

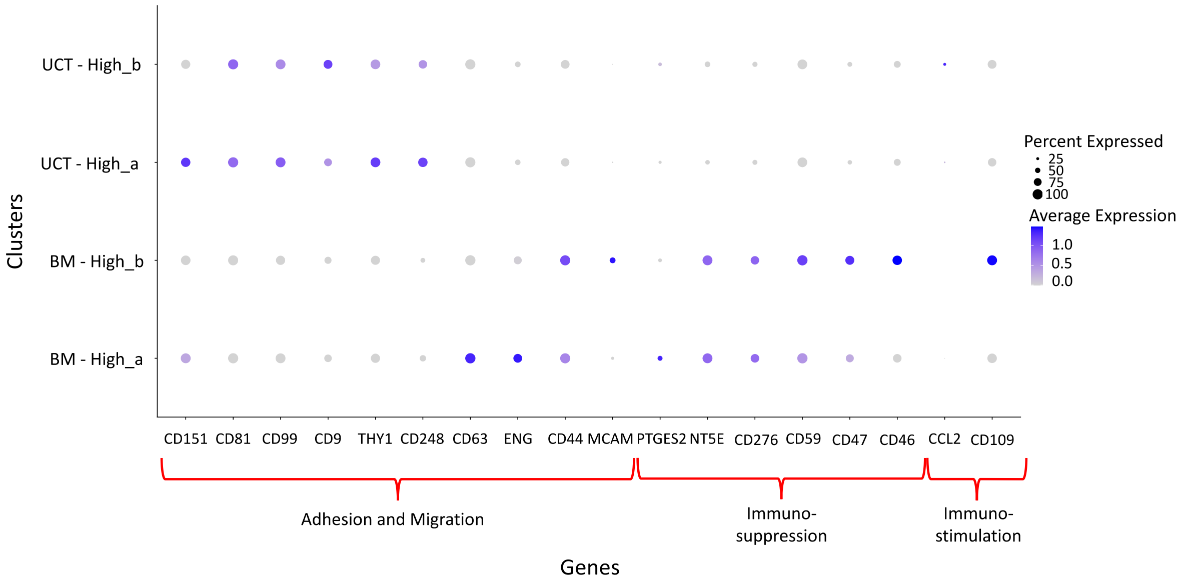


Figure S5: Dot plot displaying the average expression for cell adhesion and migration and immunomodulatory function-associated genes. The colors represent the average expression of the genes per cell. The scale is from low expression of 0 (gray) to high expression of 1.5 log counts per million (blue). The size of the dots represents the proportion of cells in each cluster that express the gene.

A.

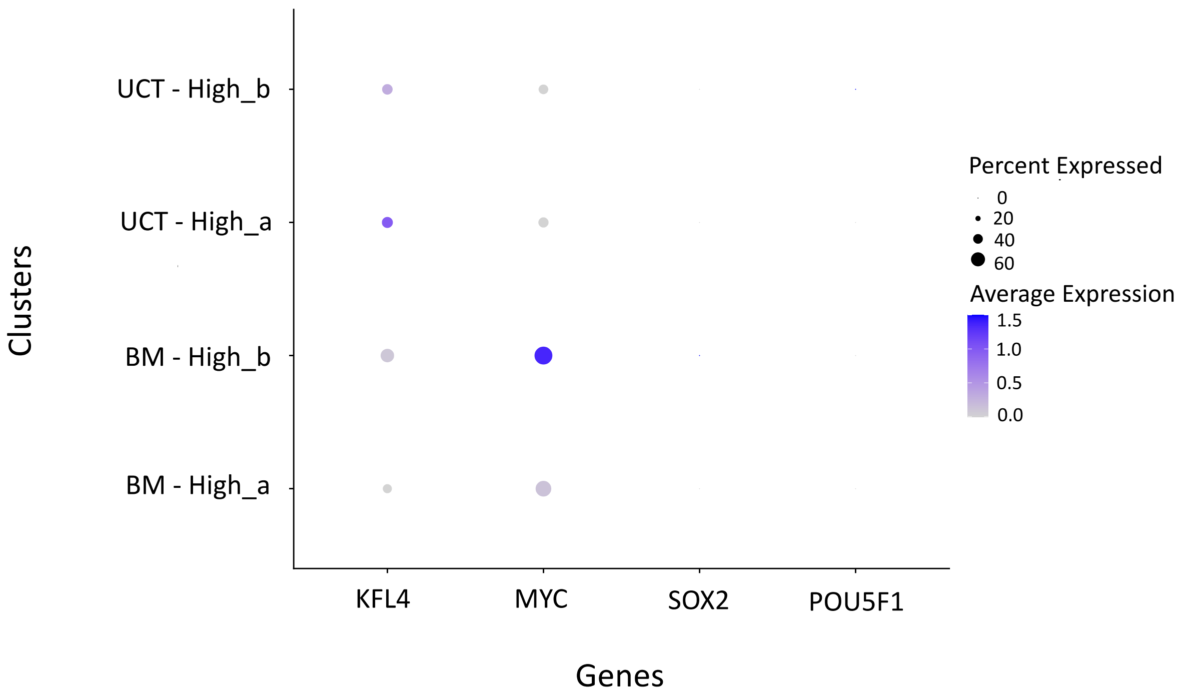


B.

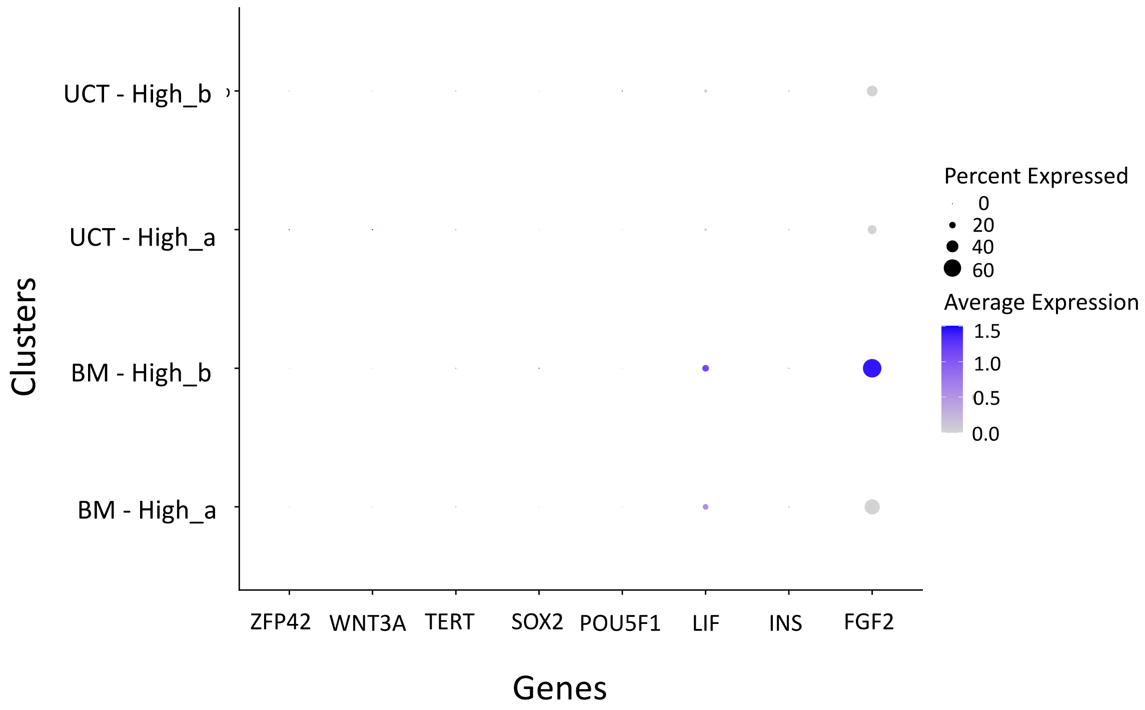


Figure S6: Dot plots displaying genes of interest. (A) This dot plot shows some pluripotent markers. (B) This dot plot displays stemness markers. The expression of some of these genes is low. There is not significant difference in the levels of expression between the clusters of MSCs.
